# Microcystin-LR aerosol exposure increases inflammatory drivers of asthma, Evidence of an NF-κB amplification mechanism

**DOI:** 10.1101/2025.09.13.676033

**Authors:** Joshua D. Breidenbach, Bivek Timalsina, Benjamin W. French, Lauren Stanoszek, John-Paul Lavik, Irum S. Breidenbach, Upasana Shrestha, Andrew Kleinhenz, Apurva Lad, Prabhatchandra Dube, Shungang Zhang, Dhilhani Faleel, R. Mark Wooten, James C. Willey, Jeffrey R. Hammersley, Nikolai N. Modyanov, Deepak Malhotra, Lance D. Dworkin, Steven T. Haller, David J. Kennedy

## Abstract

Microcystin-LR (MC-LR) is one of a large family of cyanotoxins which are naturally produced by cyanobacteria within harmful algal blooms occurring in bodies of water globally. Early findings of the toxicity of such blooms stemmed from fatalities of livestock drinking from affected water. Since then, various toxins have been identified such as the microcystins. Microcystin-LR has been studied as a representative congener due to its abundance and toxicity. While there have been extensive studies of microcystin-LR by oral route exposure, we have turned our attention to inhalation route exposure due to the recent findings of microcystin-containing lake and sea-spray aerosol. We have shown inflammatory outcomes in the airways of mice and in human cell culture models after microcystin aerosol exposure, and have found a consistent molecular patterns similar to those of Type 1/Type 17 driven neutrophilic asthma. Here we address the hypothesis that MC-LR will increase the inflammatory mediators of neutrophilic asthma leading to worsening symptoms. This is tested and characterized in both *in vitro* and *in vivo* models. We found that asthma symptoms and molecular signatures of inflammation are both worsened by MC-LR exposure in a mouse model of neutrophilic asthma. We found that 3D human airway cell culture models reconstructed from asthmatic donor cells are similarly affected, however healthy donor cells are nearly unaltered by comparison. Aggregating these findings with RNA sequencing data from all models, we developed a hypothetical molecular mechanism which relies on MC-LR mediated amplification of existing inflammatory signaling. We test this in a human reporter cell line of NF-κB activity and further demonstrate the mechanism by inhibitor testing. This study sheds light on the risk to asthmatic patients living near or recreating on affected bodies of water. Beyond asthma, we believe this study provides crucial insight into the findings over the last 40 years concerning disparate outcomes of MC-LR exposure as the result of exposure will be dependent on the signaling state of the tissue upon exposure.

## 1. Introduction

Despite the common name “blue-green algae,” cyanobacteria are an ancient phylum of Gram-negative bacteria. These photosynthetic organisms are commonly associated with the Great Oxidation Event, or the sudden oxygenation of the Earth’s atmosphere [1]; however, they continue to have a substantial impact on the environment and human health today in the form of cyanobacterial harmful algal blooms (cHABs). These overgrowth events have received attention since the 1980s, when reports of domesticated animals and livestock dying suddenly after drinking from bodies of water affected by these blooms began to emerge [2]. Today, cHABs are known to produce dermatoxins, neurotoxins, and hepatotoxins, including over 270 congeners of hepatotoxic microcystins [3].

Microcystins are cyclic peptides and are considered the most common cyanobacterial toxin. Among the microcystins, microcystin-LR (MC-LR) is the most toxic congener (LD_50_ by ingestion: 10.9 mg/kg). Named for its distinguishing leucine (L) and arginine (R) residues, MC-LR has been detected in natural water bodies around the world at levels as high as 36.5 µg/mL (∼36.6 µM) and is expected to become increasingly prevalent in the wake of changing weather patterns and anthropogenic factors [4–6]. Microcystins are traditionally considered hepatotoxins; oral exposure to MC-LR is associated with hemorrhaging of the liver and has been since early accidental exposures to whole cHAB-contaminated water [7–9]. More recently, data suggests additional systems are impacted by MC-LR including inflammatory outcomes in the lungs and gastrointestinal system as well as on immune cells directly (36137425, 31242640, 39591225, 34576099, 35330169).

MCLR promotes the expression of L-selectin, beta2-integrin, and cytokine-induced neutrophil chemoattractant-2alphabeta (CINC-2alphabeta) in neutrophils, leading to-increased chemoattraction toward fMLP [10–13]. There is also evidence of MC-LR-mediated ROS production in neutrophils [12]. Our lab previously reported that MC-LR induces the expression of the pro-inflammatory genes *Tnf* and *Il1b* in mouse and rat macrophages [14]. In lymphocytes, MC-LR induces the expression of pro-inflammatory cytokines and causes apoptosis of T and B cells [15, 16]. And in the human Jurkat T cell line, MC-LR was activates CaM kinase II and IV, which may go on to activate NF-κB via the phosphorylation of IκB or calcineurin (PP2B), leading to activation of NFAT-family transcription factors [17]. Together, these data indicate that the impact of MC-LR on the innate immune system plays a critical role in the adverse health outcomes observed after exposure to this cHAB toxin.

Beyond oral exposure, there is evidence that sea and lake spray aerosols can carry particulates over 30 kilometers inland and that MC-LR possesses molecular characteristics that support efficient aerosolization indicating that inhalation is an exposure route that warrants concern [18–20]. Notably, even systemic exposure to MC-LR is associated with increased granulocytes and the appearance of lesions in the lungs of mice [21–23]. Still, the cell signaling responses and health outcomes of inhaled MC-LR have only recently been studied *in vivo*, resulting in inflammation (39591225). Importantly, this inflammation is consistent with the inflammation of Type 1/Type 17 driven neutrophilic asthma [24]. The aim of this study was to examine the impact of inhaled MC-LR in the context of pre-existing inflammatory signaling. Herein, we demonstrate that MC-LR amplifies pre-existing inflammatory signaling in a mouse model of neutrophilic asthma. Using 3D models of human airway epithelia from both healthy and asthmatic donors, we provide evidence that MC-LR-mediated pro-inflammatory responses may originate from the epithelium rather than recruited immune cells and that this signaling proceeds in an NF-κB-mediated fashion.

## 2. Materials and Methods

### 2.1. Animals, asthma model and aerosol exposures

Male C57BL/6J (C57BL/6) mice (000664; The Jackson Laboratory, ME, USA) were 10-11 weeks old at the beginning of this study. All mice were caged socially in microisolator cages, given access to food and water *ad libitum* and kept on a 12:12-hour dark-light cycle. Protocols for animal experimentation were approved by The University of Toledo Institutional Animal Care and Use Committee (IACUC protocol #108663, approval date 9 February 2016).

For the development of neutrophilic asthma in mice, a well-reported and repeatable model was adapted to accommodate the microcystin-LR exposures (Figure 1) [25]. Briefly, mice were sensitized to a mixture of 25 µg house dust mite (HDM) + 10 µg lipopolysaccharide (LPS) in a volume of 30 µL of normal saline (0.9% NaCl) by intranasal instillation (i.n.) while under isoflurane anesthesia on days 0, 1 and 2. Beginning on day 7, mice were challenged against antigen once a day, 3 days per week, for 4 weeks. The challenge consisted of HDM by i.n. (25 µg in 30 µL of normal saline) while under isoflurane anesthesia. MC-LR was purchased from Cayman Chemicals (Item No. 10007188, Ann Arbor, MI, USA). House dust mite (*Dermatophagoides Pteronyssinus*) whole body extract was purchased from Stallergenes-Greer (Item: XPB82D3A2.5, Switzerland) and the total dry weight was used to make solutions for these experiments. LPS was purchased from Invivogen (Item: vac-3pelps, San Diego, CA, USA).

**Figure 1:**
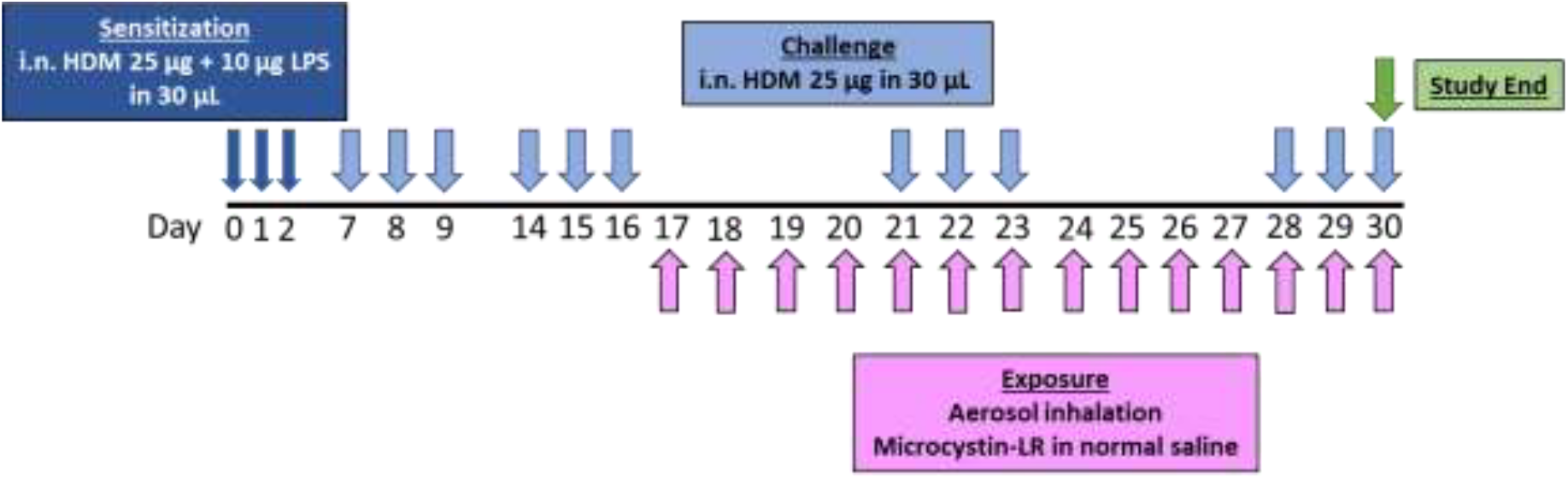
Overview of murine neutrophilic asthma model and MC-LR exposures.

Beginning at day 17 of the study, mice were exposed to MC-LR aerosol or normal saline vehicle aerosol using an inExpose (SCIREQ; Montréal, Canada) animal exposure system equipped with a nose-only adapter and a small particle (2.5-4 µm) VMD Aeroneb Lab ultrasonic nebulizer. Exposures were 1 hour each day for 14 days. On day 30, tissues were harvested within 1 hour after the final challenge and exposure. Lung deposition of 13 µg/m2 per day was chosen to mimic mass deposition of MC-LR in our previous *in vitro* study [26]. The concentration of solution required to achieve the mass deposited was related by the following equation:

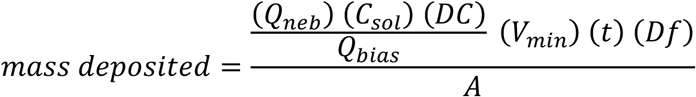

Where,

Qneb = output rate of nebulizer = 0.3 mL/min

Csol = concentration of MC-LR in solution

DC = duty cycle = 50%

Qbias = bias flow = 3.5 L/min

Vmin = minute volume of mouse = 0.04 L/min

t = exposure time = 60 min

Df = deposition fraction = 0.2

A = surface area of mouse lung = 0.02 m2 [27]

### 2.2. Histology

The left lung of each mouse was completely submerged in 10 mL of 10% buffered formalin for 24 hours on a rocker. Afterward, the formalin fixative was replaced with 70% histological grade ethanol for at least 24 hours before paraffin embedding. Whole lungs were sectioned at a thickness of 5 microns in the coronal/frontal plane and stained by hematoxylin and eosin (H&E). Images of all stained slides were collected at 20x on a VS120 Virtual Slide Microscope (Olympus, Tokyo, Japan). Histology slides were scored blindly for inflammation severity on a scale of 0-4, with 0: absent, 1: mild, 2: moderate, 3: severe, 4: very severe. Twenty random foci per section were evaluated at 20x magnification and averaged.

### 2.3. Bronchoalveolar lavage

In a separate cohort of mice than was used for histology, bronchoalveolar lavage was performed and adapted from a published method [28, 29]. Briefly, airways were lavaged with sterile normal saline (0.9% NaCl). The fluid recovered was centrifuged at 200 x g for 10 minutes, and red blood cells were lysed by 3-minute incubation with ACK lysing buffer (A1049201; Thermo Fisher; Waltham, MA, USA). After washing with 10 mL PBS, cells were resuspended and total cell counts were obtained by TC20 automated cell counter (Bio-Rad; Hercules, CA, USA). For BALF differential cell counts, slides were prepared by cytospin (2000 RPM for 3 minutes) and stained with Hema 3 Manual Staining System (22-122911; Fisher Scientific; Hampton, NH, USA). Slides were assessed by a board-certified, blinded pathologist who categorized each by cell type based on various features including size, shape and nuclear/cytoplasmic characteristics. For non-asthmatic subjects, 200 cells per slide were assessed. For asthmatic subjects (more cellular), 400 cells per slide were assessed. All cell counts were normalized by the recovered BALF volume of each animal.

### 2.4. Tissue protein measurements

Lung lobe sections weighing at least 20 mg were placed in 2 mL round-bottom centrifuge tubes with 1 mL DMEM cell medium (DML28; Caisson Laboratories, Inc., Smithfield, UT, USA) along with a stainless steel bead. Tissues were mechanically lysed using a TissueLyser II (QIAGEN; Hilden, Germany) at 25 Hz for 10 minutes. Protein measurement services were performed by Quansys Biosciences (Logan, UT, USA). Custom multiplex ELISA-based Q-Plex™ technology delivered quantification of murine protein cytokines and chemokines: Il-1α, Il-1β, Il-2, Il-3, Il-4, Il-5, Il-6, Il-10, Il-12p70, Il-13, Il-17A, Mcp-1 (Ccl2), Ifna, Ifnb, Ifnγ, Tnfα, Mip-1α (Ccl3), Gm-csf, Rantes (Ccl5), Eotaxin (Ccl11), Mip-2 (Cxcl2), Kc (Cxcl1), MDC (Ccl22), TARC (Ccl17), TCA-3 (Ccl1). For results of all analytes, please see Table C.1. All protein concentrations were calculated as pg analyte per mg total protein.

### 2.5. 3D human primary airway epithelium model and exposures

A 3D model of cultured primary human airway epithelial cells (MucilAir™; Epithelix; Geneva, Switzerland) was established comprising cells pooled from 14 healthy donors or an asthmatic donor. Cell cultures were successfully established on 24-well transwell inserts at an air-liquid interface with serum-free MucilAir™ Medium (Epithelix; Geneva, Switzerland). All treatment groups were completed in triplicate.

To expose cell cultures to test materials, saline solutions containing MC-LR (GreenWater Laboratories; Palatka, FL) were nebulized using a Vitrocell-Cloud chamber which is designed to deliver dose-controlled and spatially uniform deposition of liquid aerosols to the apical side of cells cultured at the air-liquid interface. Exposures occurred once per day for 3 days, and during each of the 3 exposures, the aerosol was generated and deposited to the apical surface within approximately 3 minutes. 3 minutes of aerosol generation was necessary for uniform aerosol deposition. The nebulizer can deliver 200 µL of liquid per exposure and the bottom surface of the Vitrocell-Cloud chamber is 136.5 cm2 while the inserts are 0.33 cm2 each. Assuming a 90% delivery efficacy, this equals approximately 0.43 µL of deposition per insert following exposure by this system. Therefore, the 1 µM MC-LR saline solution yielded an approximate deposition of 12900 ng/m2.

### 2.6. Functional Tissue Assays

Functional tissue assays were performed in the 3D airway epithelium model as described previously [26]. Briefly, cytotoxicity and tissue integrity were assessed by lactate dehydrogenous (LDH) and transepithelial electrical resistance (TEER) assays, respectively. LDH was measured in the conditioned media of the MC-LR exposed cell models reconstructed from healthy and asthmatic donor cells. To determine the percentage of cytotocicity, the following equation was used: Cytotoxicity (X) = (A (exp value) –A (low control)/A (high control) – A (low control)) X 100. Where A = absorbance values and the high control value was obtained by 10 % Triton X-100 apical treatment. In the TEER assay, an electrode is placed at the apical and basal sides of the epithelial tissue inserts and the resistance to electric current is measured. A loss of resistance indicates a loss of physical tissue integrity.

### 2.7. Gene Expression

Gene expression of mouse lung tissue began with RNA extraction from snap frozen tissue samples. RNA was isolated utilizing the Qiagen RNeasy Plus Mini Kit (Qiagen, Germantown, MD, USA, Catalog No. 74134) and the Qiagen QIAcube extraction methodology. Gene expression of airway epithelial cell inserts began with RNA extraction from snap frozen tissue inserts. RNA was isolated utilizing the Qiagen RNeasy Plus Mini Kit with slight modifications to the protocol, as published [26]. Concentration and purity values were determined by NanoDrop (Thermo Fisher Scientific, ND-1000; Waltham, MA) and were consistently adequate for analysis (260/280 > 1.8, 260/230 > 1.0, RNA amount > 2 µg).

For transcriptomic analysis of the murine RNA, RNA sequencing with PolyA selection was performed by Azenta Life Science (formerly Genewiz) (Plainfield, NJ, USA). Prior to sequencing, concentration and purity values of the samples were confirmed. Reads were uploaded to the Galaxy web platform using a public server at usegalaxy.eu to analyze the data [30]. Reads with adapter contamination, low quality bases, and undetermined bases were removed using Cutadapt, and remaining reads were mapped to the mouse genome (Mus Musculus, mm10 full) using HISAT2 [31, 32]. Transcripts were assembed using featureCounts [33].

For transcriptomic analysis of the human primary airway epithelium model, Poly(A) RNA sequencing was performed by LC Sciences (Houston, TX). Prior to sequencing, concentration and purity values of the samples were confirmed. The Poly(A) RNA sequencing library was prepared following Illumina’s TruSeq-stranded-mRNA sample prepartion protocol and paired-ended sequencing was performed on Illumina’s NovaSeq 6000 (San Diego, CA). Quality control statistics were generated using FastQC. Reads with adapter contamination, low quality bases, and undetermined bases were removed using Cutadapt, and remaining reads were mapped to the human genome (Genome Reference Consortium Human Build 38 patch release 13 (GRCh38.p13); GCA_000001405.28) using HISAT2 [31, 32]. Transcripts were assembled using StringTie [34].

In both cases, counts tables were analyzed using the edgeR package on the Galaxy Platform (https://usegalaxy.eu/) and differential expression of genes were calculated, resulting in log base 2 of the fold change (log2FC) values [30, 35]. When necessary, gene IDs were converted using GeneToList (https://www.genetolist.com/) [36]. Through edgeR, a weighted trimmed (T) mean (M) of the log expression ratios (M) (TMM) based method was used to generate normalized counts as counts per million (CPM) for graphical representation [35, 37]. Only genes with CPM values greater than 0.5 in at least half of the samples were considered for the analysis. All pathway analyses were performed with the web tool suite g:Profiler using the g:GOSt functional profiling application [38].

### 2.8. Transcription signature comparisons

For the comparision of the asthma expression signature in the human primary epithelial cells with published datasets, we utilized the Library of Integrated Network-Based Cellular Signatures (LINCS), a large multi-omics profiling database. Integrative LINCS (iLINCS) is a web platform for the analysis of LINCS datasets that was developed under the LINCS consortium (www.ilincs.org/). iLINCS uses weighted Pearson correlation analysis to measure the concordance between signatures [39].

### 2.9. Protein-protein interactions

The online web tool suite STRING was used to assess protein-protein interactions using a database of both predicted and experimentally-determined protein interactions [40]. Proteins were entered as their respective gene names in the “Multiple proteins” tool and *Homo sapiens* was selected. Specific settings used were: Network type – full, Minimum confidence – medium, 1st shell maximum – 10, 2nd shell maximum – 30, Meaning of network edges – confidence, Active interaction sources – all.

### 2.10. Reporter cell line

The A549-Dual reporter cell line was purchased from Invivogen (Item: a549d-nfis, San Diego, CA, USA). Cells were handled according to the commercial guidelines. Growth medium was phenol red free DMEM with 10% heat-inactivated FBS (10438026; Thermofisher), 100 µg/mL pen-strep (P0781; Sigma-Aldrich), 100 µg/mL normocin (ant-nr-1, Invivogen), 10 µg/mL blasticidin (ant-bl-05; Invivogen), and 100 µg/mL zeocin (ant-zn-05; Invivogen). NF-κB activity assays were performed via the QUANTI Blue Solution (rep-qbs; Invivogen) assay, following the manufacture’s protocols in a 96-well plate format. 20 µL of MC-LR solution or water vehicle were added to an empty 96-well plate. 180 µL of cell suspension was added to each well for ∼50,000 cells/well. Plates were incubated at 37°C in a CO2 incubator for 24 hours. Then, plates were gently washed with PBS twice before resupplying 180 µL of fresh medium. Next, 20 µL of each exposure agent (IL1β, MC-LR, or vehicle) were added to the wells. Plates were again incubated at 37°C + CO2, and 20 µL of the conditioned media was sampled at the time points shown. This conditioned media was combined with 180 µL of assay reagent and incubated for approximately 3 hours at 37°C before absorbance measurements were taken at OD655.

For the testing of the PS-1145 inhibitor, a similar pre-treatment and exposure process was performed. 20 µL of MC-LR solution, PS-1145 solution, MC-LR solution with PS-1145 solution or vehicle (0.5% DMSO in water) were added to an empty 96-well plate. 180 µL of cell suspension was added to each well for ∼50,000 cells/well. Plates were incubated at 37°C in a CO2 incubator for 24 hours. Then, plates were gently washed with PBS twice before resupplying 180 µL of fresh medium. Next, 20 µL of each exposure agent (IL1β, MC-LR, PS-1145 or vehicle) were added to the wells. Plates were again incubated at 37°C + CO2, and 20 µL of the conditioned media was sampled after 24 hours.

MC-LR was sourced from Cayman Chemical (Item no. 10007188; Ann Arbor, MI, USA) and used at a final concentration of 1 µM. PS-1145 was also sourced from Cayman Chemical (Item no. 14862) and used at a concentration of 20 µM. Human recombinant IL-1β was sourced from Invivogen (Item. rcyec-hil1b) and used at a final concentration of 0.5 ng/mL.

### 2.11. Statistics

The statistical tests in this work were completed in GraphPad Prism version 7.0.5 for Windows (GraphPad Software; San Diego, California, USA, https://www.graphpad.com). Statistics by Student’s t-test between each group and healthy vehicle, and between MC-LR-exposed and respective vehicle controls. *, **, ***, **** indicates p ≤ 0.05, 0.01, 0.001, 0.0001, respectively.

## 3. Results

### 3.1. Increased Immune Cell Infiltration in Asthmatic MC-LR-Exposed Mice

To understand the risk of adverse consequences to patients with pre-existing airway inflammation from MC-LR inhalation exposure, we modeled neutrophilic asthma in mice. Male C57BL/6 mice were sensitized to a mixture of HDM and LPS and then challenged with HDM 3 days per week for 4 weeks (Figure 1). After 2 weeks, mice were exposed to MC-LR aerosol or vehicle by nose-only inhalation 1 hour a day for 14 days (Figure 1). The estimated deposition of MC-LR to the lungs each day was 13 µg/m2 (see Methods section 2.1). Lung tissues were prepared for histology and molecular evaluation within 1 hour after the final exposure.

Lung sections were stained by H&E and scored for inflammation severity (Figure 2). Consistent with our previous studies, MC-LR exposure led to a significant increase in inflammation in the lung tissue of healthy mice. As expected, increases were also found in the asthmatic control mice compared with healthy controls. No further increases were observed in MC-LR-exposed asthmatic mice, although it was the group with the greatest significance compared with vehicle.

**Figure 2:**
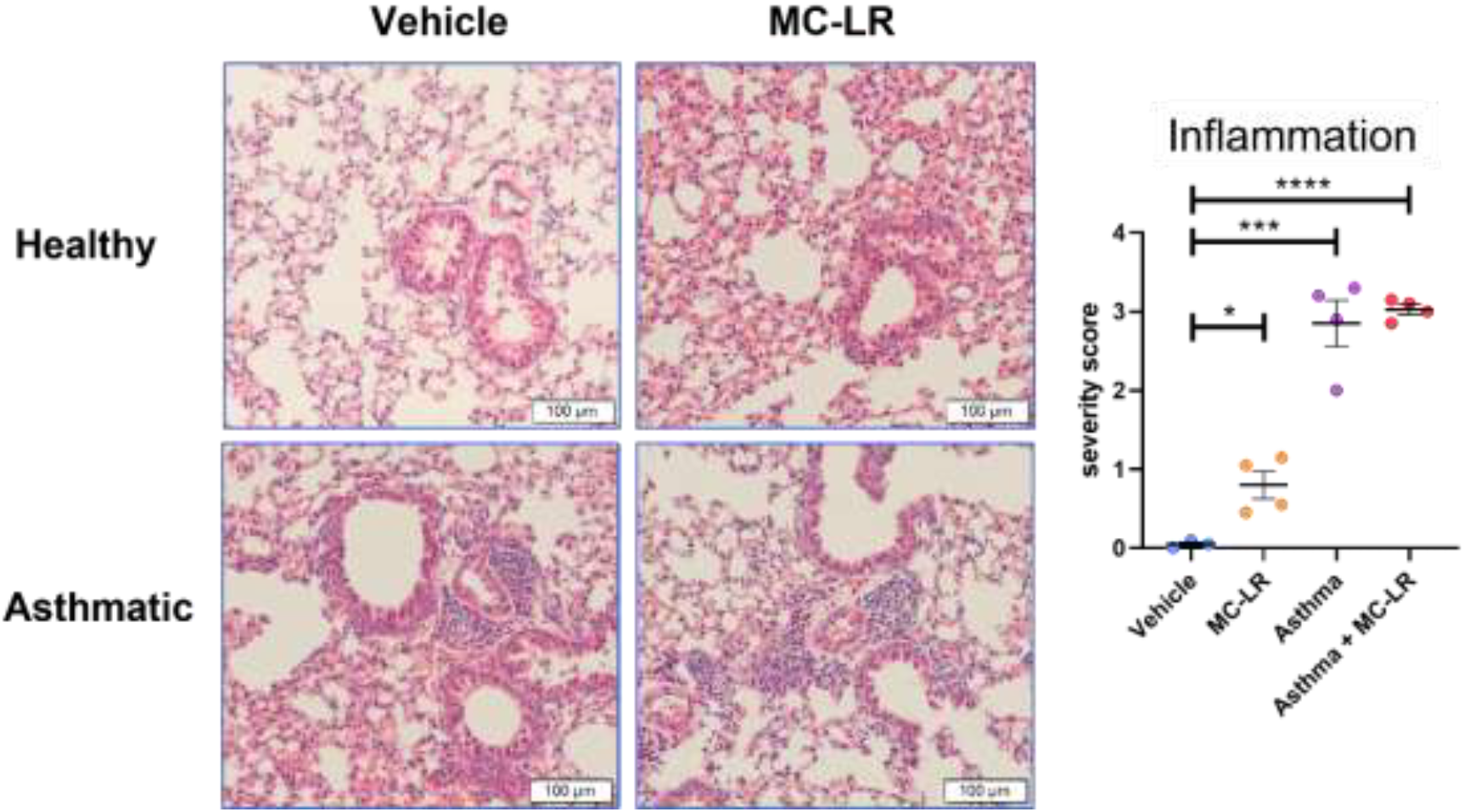
Histological evaluation of the lung. A) Representative images of H&E-stained sections of healthy or asthmatic mouse lung after the complete MC-LR or vehicle aerosol exposure regimen. Severity scoring of inflammation in the H&E-stained sections.

To further evaluate immune infiltrates of the airways, bronchoalveolar lavage was performed on a separate set of mice (Figure 3). As previously reported, MC-LR exposure led to increases in the total cells in the BAL which consisted of increased neutrophils and macrophages. When compared with healthy mice, increased neutrophils and eosinophils and a positive trend for lymphocytes was observed in asthmatic mice following MC-LR exposure. Importantly, asthmatic mice demonstrated further increases of total BAL cells, eosinophils, and lymphocytes, with a potential trend in neutrophils following MC-LR exposure when compared with all other groups.

**Figure 3:**
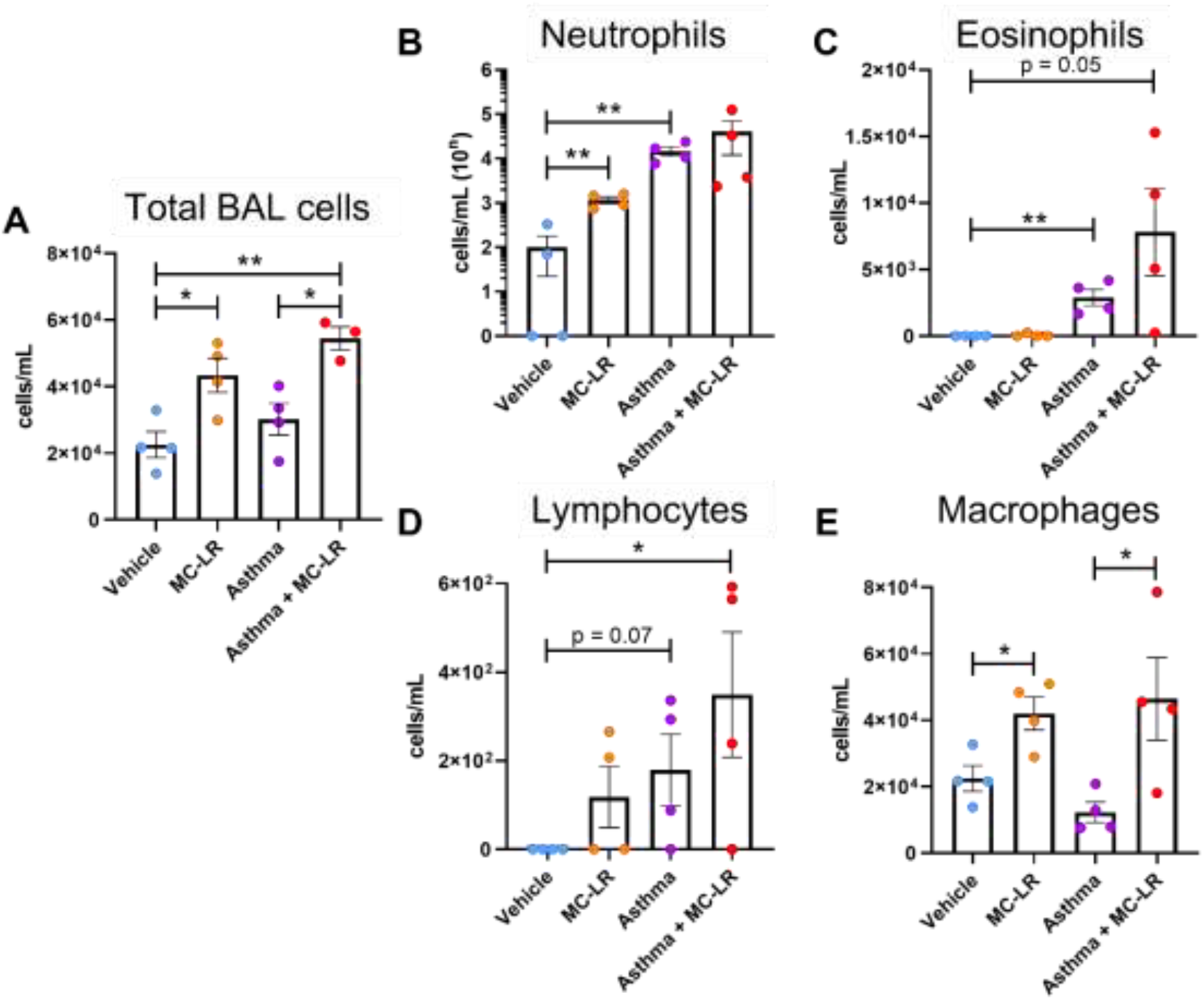
Bronchoalveolar lavage of asthmatic and healthy mice exposed to MC-LR or vehicle. Cellular constituents of BAL fluid normalized to the recovered volume. A) total cells, B) neutrophils, eosinophils, D) lymphocytes, and E) macrophages. *, ** indicates p ≤ 0.05, 0.01, respectively. Statistics by Student’s t-test between each group and vehicle, and between MC-LR-exposed and respective vehicle controls.

### 3.2. Increased markers of inflammation in asthmatic MC-LR exposed mice

Markers of inflammation including cytokine and chemokine proteins were evaluated from lung lysate (Figure 4). Similar to previous observations, MC-LR aerosol exposure in healthy mice led to increased type 1 and type 17 markers (Il-2 and Il-17a) (Figure 4A) as well as mixed granulocytic mediators (Cxcl1, Ccl3, Ccl22, and Ccl1) (Figure 4C). Increases were not found for type 2 markers (Il-4, Il-5, and Il-13) (Figure 4B). As expected, the neutrophilic asthma model displayed significant increases in type 1/type 17 and the mixed granulocytic, but not the type 2 markers (Figure 4). Importantly, MC-LR exposure in asthmatic mice led to further significance in Cxcl1, Ccl22, and Ccl1, and significant increases in Mip-1a (Ccl1) and Il-1a, with trends in Il-2 and Il-17a (Figure 4A and C). In the type 2 markers in asthmatic mice, MC-LR exposure brought about significant decreases in Il-13 and decreasing trends in Il-4 and Il-5 (Figure 4B).

**Figure 4:**
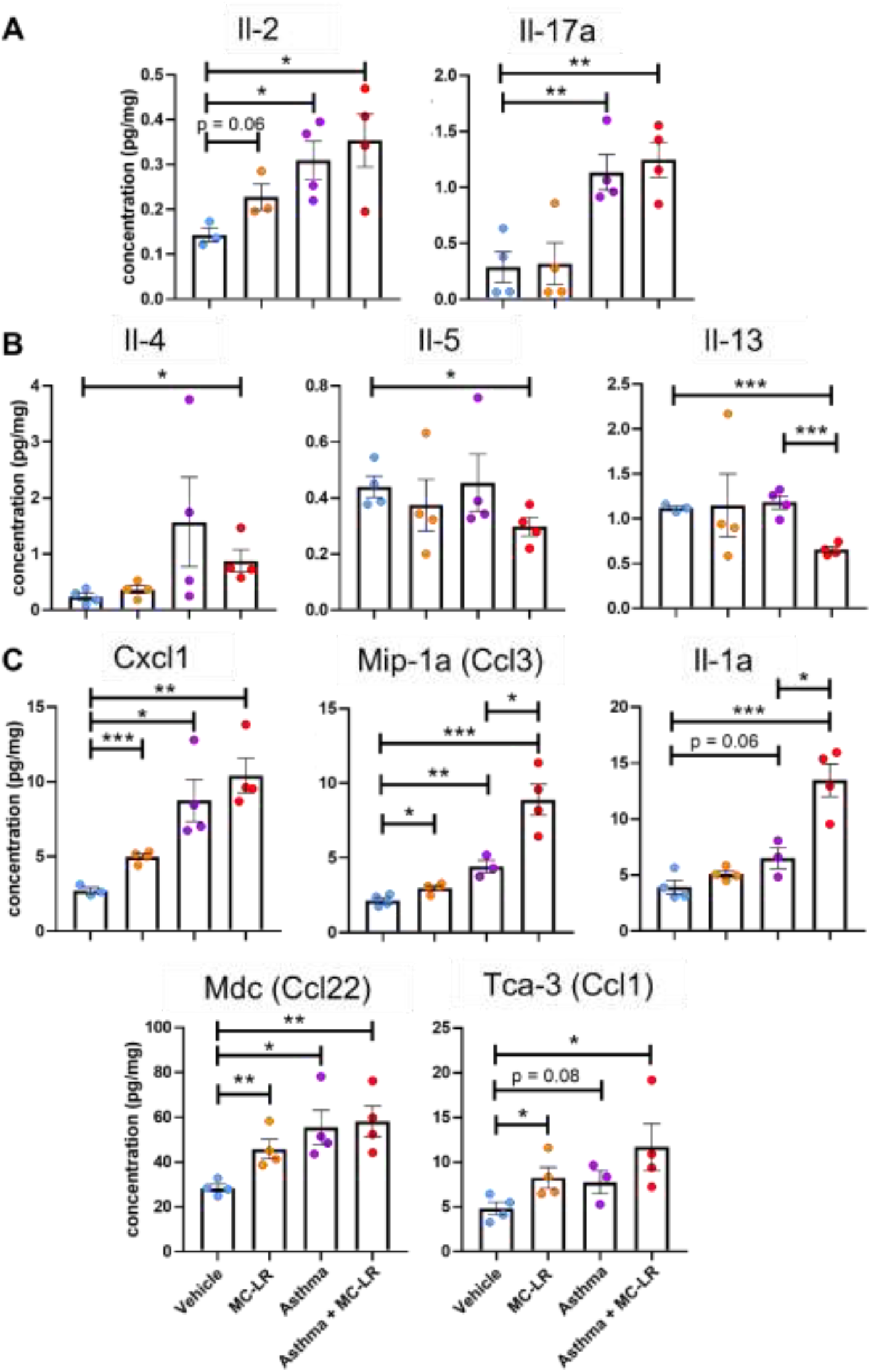
Concentration of type 1/type 17 cytokine and chemokine proteins from lung lysates of MC-LR or vehicle exposed healthy or asthmatic mice. *, **, *** indicates p ≤ 0.05, 0.01, 0.001, respectively. Statistics by Student’s t-test between each group and vehicle, and between MC-LR exposed and respective vehicle controls.

Lung lysates were also utilized for gene expression analysis by RNA-sequencing. In our analysis, genes of interest were screened by searching for those with increased expression (> 0.5 log2FC) between the asthmatic controls and the healthy vehicle controls, and then further increases (> 0.5 log2FC) in the MC-LR exposed asthmatics compared with the asthmatic controls. In this way, protein coding genes upregulated in asthma, with MC-LR-mediated exacerbated expression were selected. Gene ontology biological process (GO-BP) pathway enrichment analysis of these genes revealed associations with “neutrophil migration”, “granulocytic migration”, “chemokine-mediated signaling pathway”, and others (Figure 5A). Additional pathway enrichment via the Kyoto Encyclopedia of Genes and Genomes (KEGG) resulted in associations with “rheumatoid arthritis”, “IL-17 signaling pathway”, “viral interaction with cytokine and cytokine receptor”, and others (Figure 5A). Genes found to be associated with the GO-BP “neutrophil migration” pathway were *Ccl2, Cxcl2, Cxcl3, Cxcl5, Gp2*, and *Jaml* (Figure 5B). Between MC-LR and vehicle healthy mice, we found significant upregulation of *Ccl2, Cxcl2, Cxcl5, Gp2*, with trends in *Cxcl3*, and *Jaml*. Asthmatic control mice were found to have significant increases in all of these genes besides *Cxcl2* when compared to healthy controls. These increases grew in significance in the MC-LR asthmatic mice in *Cxcl3, Cxcl5*, and *Jaml*, and became significant in *Cxcl2*. Finally, comparing the asthmatic MC-LR exposed mice to the asthmatic controls, there was a significant increase in *Cxcl5* with trends in all genes (Figure 5B).

**Figure 5:**
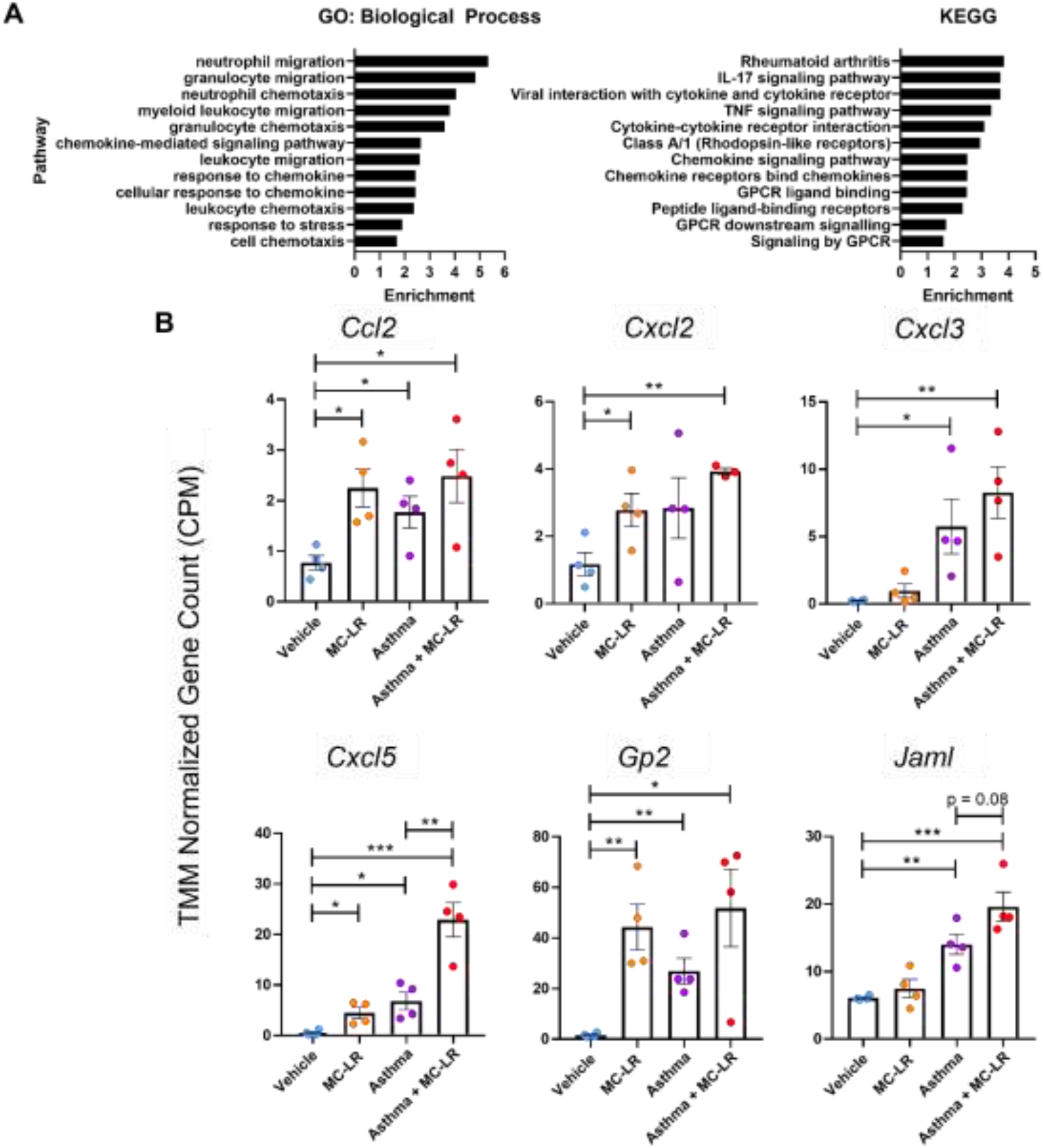
Gene expression in MC-LR- or vehicle-exposed healthy or asthmatic mice by RNA-sequencing. A) Pathway enrichment analysis of protein coding genes upregulated in asthma and exhibiting MC-LR-mediated exacerbation. B) Transcriptional levels of genes associated with the Gene Ontology Biological Process pathway “neutrophil migration.” Displayed as TMM normalized CPM values. *, **, ***, indicates p ≤ 0.05, 0.01, 0.001, respectively. Statistics by Student’s t-test between each group and vehicle, and between MC-LR exposed and respective vehicle controls.

### 3.3. MC-LR exacerbates asthmatic expression profile in human airway epithelium

To understand if the airway epithelium may be responsible for the MC-LR-mediated inflammation, we utilized a 3D model of primary human airway epithelium reconstructed from cells derived from healthy and asthmatic donors. Airway epithelium inserts were exposed to MC-LR or vehicle saline by aerosol deposition on the apical side of an air-liquid interface culture system (Figure 6A). The concentration of MC-LR solution aerosolized was 1 µM MC-LR which yielded an approximate deposition similar to the *in vivo* exposures (∼13 µg/m2) per day, for 3 days. Tissue functional measurements of cytotoxicity and tissue integrity were collected. However, no deleterious consequences were detected (data not shown).

**Figure 6:**
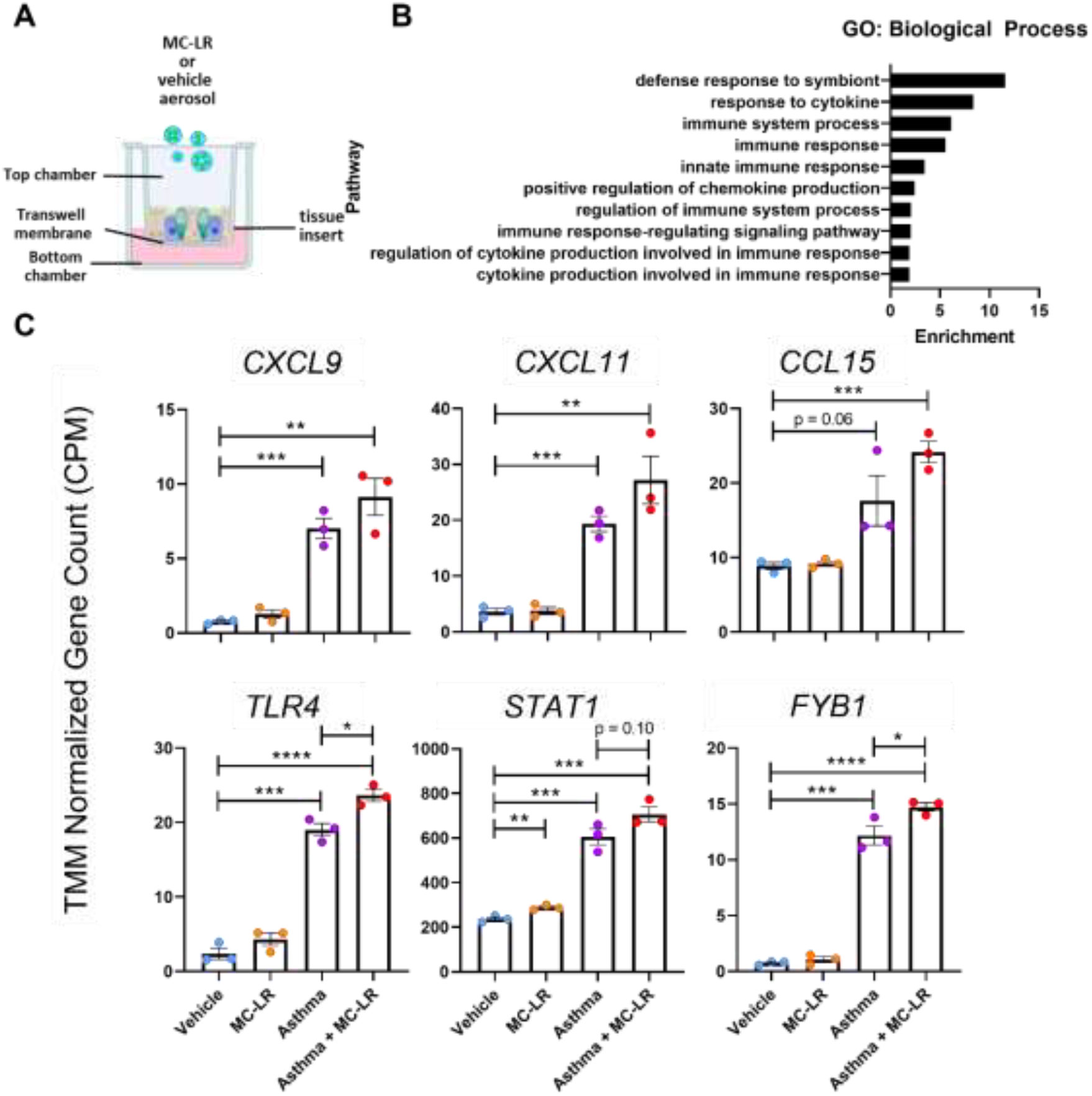
Gene expression analysis of MC-LR-exposed 3D cultured primary human airway epithelium reconstructed from cells derived from healthy and asthmatic donors. A) Culture model. B) Pathway enrichment analysis of protein coding genes upregulated in asthma and exhibiting MC-LR mediated exacerbation. C) Transcriptional levels of select genes associated with the shown pathways. Displayed as TMM normalized CPM values. *, **, ***, **** indicates p ≤ 0.05, 0.01, 0.001, 0.0001, respectively. Statistics by Student’s t-test between each group and vehicle, and between MC-LR exposed and respective vehicle controls.

Transcriptional changes induced by the donor disease status, or the MC-LR exposure were evaluated in the tissue inserts frozen 24 hours after the third and final exposure. Differential expression by RNA sequencing in the asthmatic donor cells compared with the healthy donor cells resulted in 4122 upregulated (log2FC > 0.2) protein coding genes accounting for 29% of the total detected genes. Similarly, 3887 protein coding genes were downregulated. When comparing this disease-mediated expression profile with 9097 published expression signatures using iLINCS (ilincs.org), the greatest correlation (Pearson r = .293, p = 4.8 × 10^-141) was with another differential expression signature (signature: GDS_8731) between allergic and healthy donor derived human primary airway epithelium.

Importantly, between the MC-LR exposed asthmatics and the asthmatic controls, 474 or 11% of the asthma-upregulated genes were found to be further upregulated (log2FC > 0.2). Similarly, 11% of the asthma-downregulated genes were further downregulated. Therefore, MC-LR appeared to exacerbate the asthma-mediated gene expression profile. GO-BP pathway enrichment analysis of the MC-LR-amplified, asthma-upregulated genes revealed associations with “defense response to symbiont”, “response to cytokine”, “immune system process”, “immune response”, and others (Figure 6B). The individual transcriptional levels of select genes belonging to these pathways are shown (Figure 6C). MC-LR exposure of the healthy epithelial cells induced significant upregulation of *STAT1* and trends in *CXCL9* and *TLR4*. Compared with the healthy donor cells, asthma controls displayed significant increases in *CXCL9, CXCL11, TLR4, STAT1*, and *FYB1* with a trend in *CCL15*. Exposure of the asthmatic donor epithelial cells led to significant increases in *TLR4* and *FYB1*, with trends in *CXCL9, CXCL11, CCL15*, and *STAT1* (Figure 6C).

### 3.4. Mechanistic Insight for MC-LR amplificaiton of pro-inflammatory signaling

Because it appears that the healthy and asthmatic human airway epithelium exposed to MC-LR further expresses mediators of inflammation, we set out to further understand the underlying mechanism of this response. The known mechanisms of MC-LR toxicity involve the cellular uptake through OATPs and the inhibition of PP1 and PP2A. To elucidate a potential connection between this known mechanism and our results, we analyzed protein-protein interactions between PP2A (*PPP2RA*) and select upregulated genes *TLR4, CXCL9*, and *CXCL11* (Figure 7). A k-means clustering of these interacting proteins shows that PPP2RA interacts strongly with other PP2 family members (Light Blue). In addition, CXCL9 and CXCL11 interact strongly with other chemokines (Dark Blue). Importantly, PP2A and the chemokines connect through interactions which make up the signaling mediators between the TLRs and chemokine driving transcription factors (Green, Yellow, and Red).

**Figure 7:**
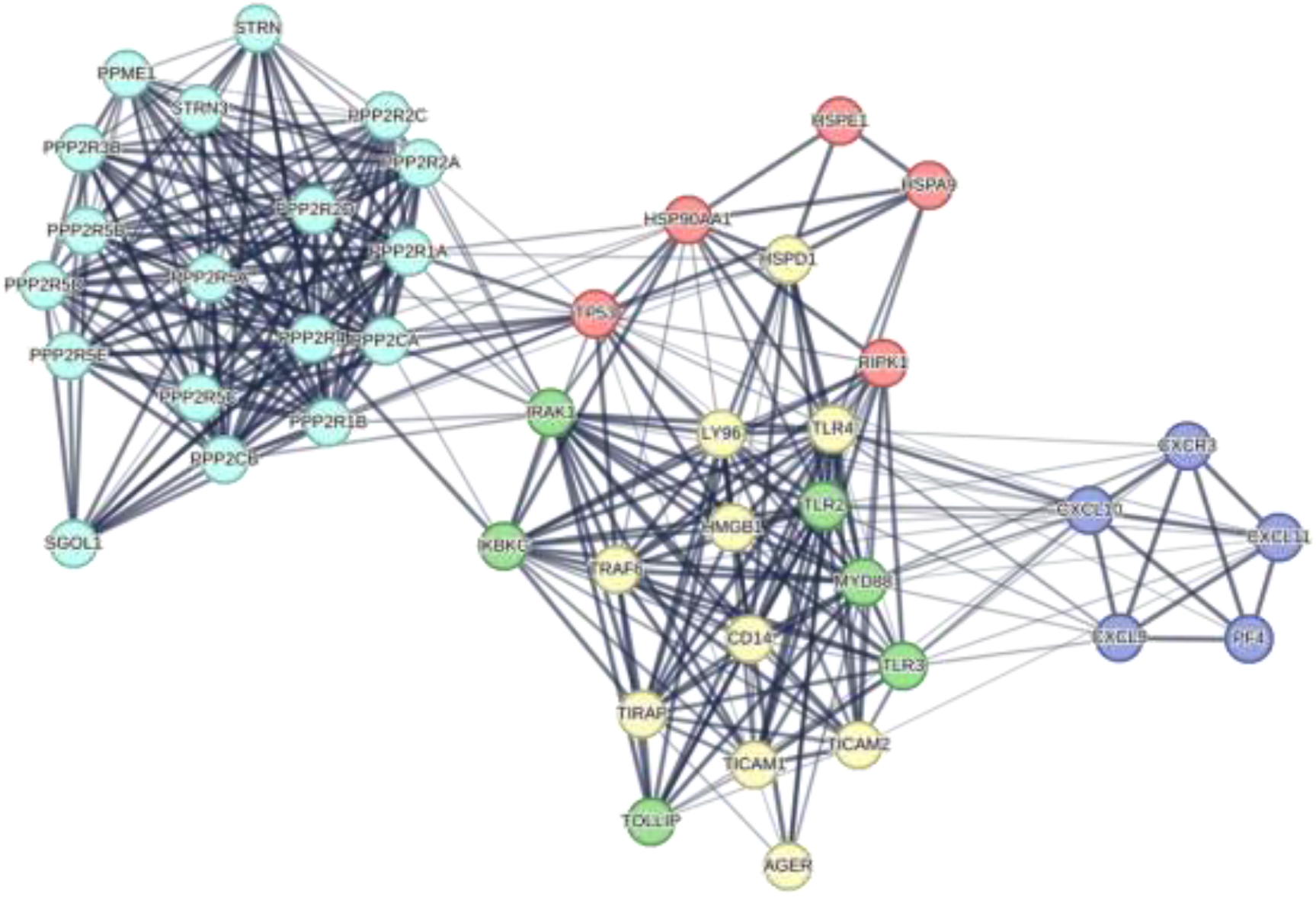
Protein-protein interaction analysis between PP2A and immune-related markers in the MC-LR exposed asthmatic human airway epithelium. K-means clustering to 5 clusters revealed groupings of PP2A-family related genes (light blue), chemokine related genes (dark blue), and signaling mediators related to both PP2A and the chemokines (green, yellow and red).

Specifically, we identified IKBKG, IRAK1, TRAF6, and MYD88, which together implicate NF-κB as an important mediator in the epithelial cell response to MC-LR.

PP2A is known to dephosphorylate IKK, acting as a suppressor for NF-κB. Therefore, we hypothesized that MC-LR functions to amplify NF-κB activity by inhibiting PP2A. To test this, we utilized A549-Dual cells, a human airway epithelial cell line which reports on NF-κB activity. We evaluated MC-LR-mediated NF-κB activity in the context of a pro-inflammatory stimuli (IL-1β) (Figure 8). Cells were pretreated for 24 hours with 1 µM MC-LR or vehicle, and then exposed to IL-1β (0.5 ng/mL), 1 µM MC-LR, or vehicle for the times shown. MC-LR pretreatment yielded no detectable change in constitutive NF-κB in the vehicle exposed cells (blue lines). However, the MC-LR exposure resulted in slight, but significant NF-κB activity, regardless of pretreatment (orange lines). As expected, IL-1β exposure led to significant increases in NF-κB activity (dashed red line). Importantly, MC-LR pretreatment was found to significantly increase the response to IL-1β (solid red line).

**Figure 8:**
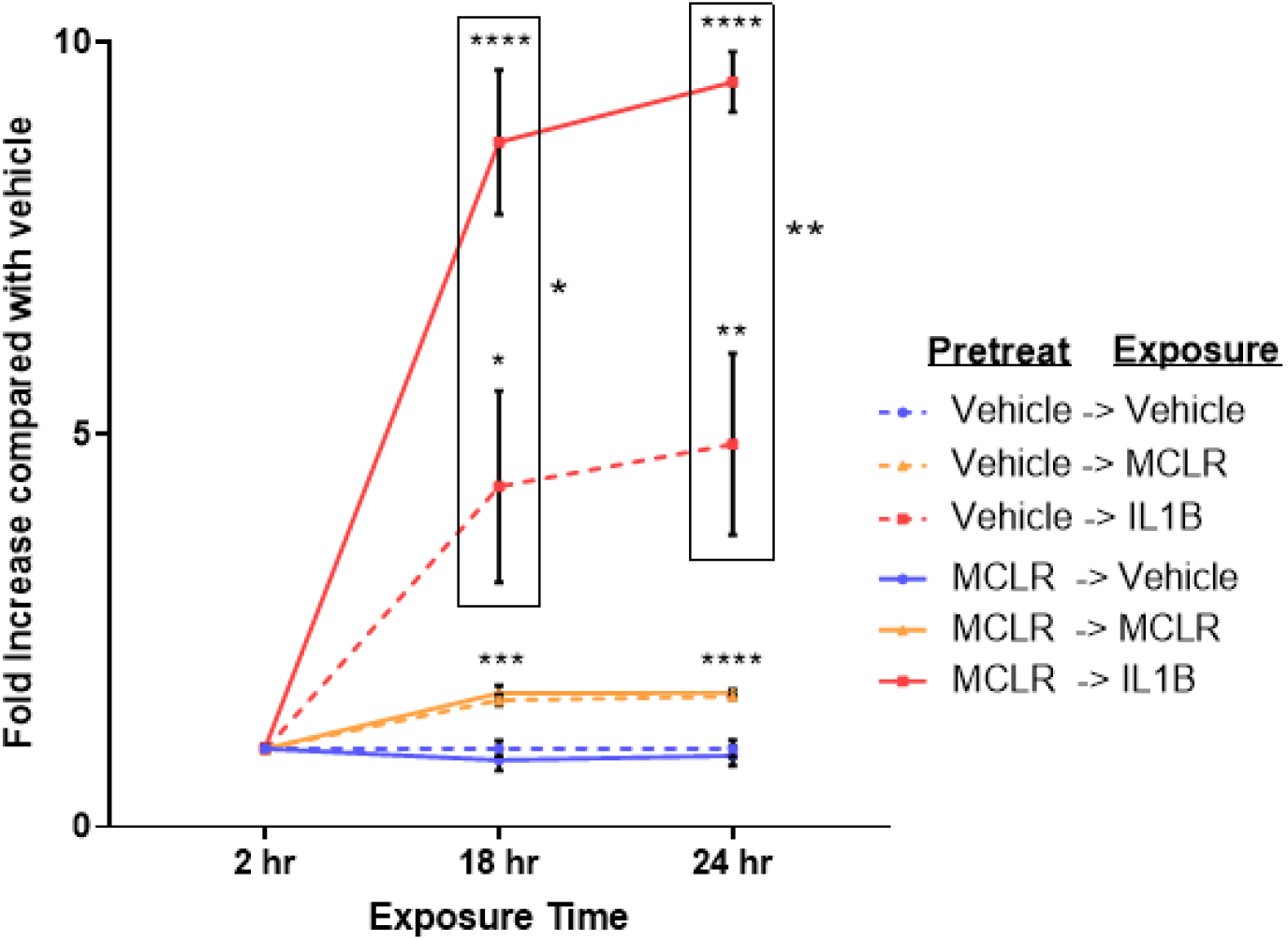
NF-κB activity measured in the A549-Dual reporter cell line. Cells were pretreated for 24 hours and exposed as indicated. *, **, ***, **** indicates p ≤ 0.05, 0.01, 0.001, 0.0001, respectively. Statistics by Student’s t-test between each group and vehicle pretreated vehicle exposed. Additional comparisons by Student’s t-test between each MC-LR pretreated and respective vehicle pretreated groups, indicated by boxed regions.

To further test, the molecular mechanism of MC-LR mediated inflammation, we performed an inhibition study. PS-1145 was chosen as it is a potent inhibitor of IKK. Therefore, it should serve to reapply the inhibitory effect of the NF-κB signaling cascade that MC-LR prevents. In the same reporter cell line used for Figure 8, we tested pre-treatment with MC-LR with and without additional pretreatment of PS-1145. Upon PS 1145 pre-treatment, we found significant inhibition of the MC-LR mediated amplification of NF-κB activity. With vehicle pretreatment, IL1B yielded 5.6 fold increase of NF-κB activity with a SEM of 0.48. With MC-LR pretreatment, IL1B yielded 19.8 fold increase of NF-κB activity with a SEM of 0.70. With MC-LR in combination with PS-1145 pretreatment, IL1B yielded 2.6 fold increase of NF-κB activity with a SEM of 0.21.

## 4. Discussion

Here for the first time, we have shown that MC-LR seems to amplify pre-existing inflammatory signaling in the airways of mice, including asthma-related signaling. Additionally, we have shown that this may begin at the epithelium and have provided evidence of MC-LR-mediated amplification of NF-κB activity in airway epithelial cells.

First, we modeled neutrophilic asthma and performed exposure experiments similar to previously completed in healthy mice [29]. Although we did not find significant increases in inflammation of the lung tissue by histology in the asthmatic MC-LR exposed mice, we did find significant increases of immune infiltrates in the BALF. Protein and gene expression analysis revealed increased type 1 and type 17 immunity markers, such as Il-2, Il-17a, Ccl3. As we have previously reported in a strain comparison study, there were little or no increased markers of type 2 immunity, such as Il-4, Il-5, or Il-13 post MC-LR exposure [29]. Genes which were upregulated in the asthmatic lung and further upregulated by MC-LR exposure associated with pathways relevant to type 1 and type 17 immunity, including “neutrophil migration” and “IL-17 signaling.” These results were consistent with findings of polymorphonuclear cells in the lungs after systemic administration of MC-LR by i.p. [23, 41–43].

To dissect the responses at the epithelium from the whole organism, we modeled human airway epithelium from healthy and asthmatic donors. There were no detectable deleterious consequences on cytotoxicity and tissue integrity. Differential gene expression by RNA sequencing revealed many genes were altered by the disease state and then further upregulated or downregulated by MC-LR exposure. Asthma-upregulated genes which were further upregulated by MC-LR exposure were associated with immune-related processes such as “defense response to symbiont” and “response to cytokine.” Importantly, these genes included the type 1 immunity driving chemokines *CXCL9* and *CXCL11* [44].

We took a hypothesis-generating bioinformatics approach to connect the outcomes in the epithelial cells to known mechanisms of PP2A inhibition [45]. We reasoned that NF-κB responses may be amplified by MC-LR exposure, and tested this possibility in an airway epithelial cell line which reports on NF-κB activity. Indeed, NF-κB activity induced by IL-1β was greatly increased after MC-LR pretreatment.

Although MC-LR mediated inflammation in the context of asthma is investigated here, the next steps our studies could explore include the evaluation of asthmatic lesions in the asthmatic mouse lung histology. This would enable us to understand if the increased inflammation leads to further deleterious changes in the tissue such as airway remodeling and excessive mucus production. Furthermore, we could evaluate differential neutrophils in the model by measuring neutrophil elastase activity.

Taken together, our results are consistent with our previously reported findings concerning the pro-inflammatory effects of MC-LR inhalation exposure. Any further research in microcystin inhalation exposure should consider patients with pre-existing inflammatory disease and the role of the epithelial cells.

## Author Contributions

JDB, BWF, RMW, JCW, JRH, NMM, STH, and DJK conceived the study. JDB, BWF, ISB, STH, and DJK wrote the manuscript. JDB, BWF, LS, JPL, ISB, AK, AL, PD, SZ, and DF performed the experiments. JDB, BWF, LS, and JPL analyzed the data. RMW, JCW, JRH, NNM, LDD, STH and DJK supervised the project. JDB, DM, STH, and DJK acquired funding for the work. All authors proofread, edited, and approved of the manuscript.

## Acknowledgments

The authors would like to thank the Women & Philanthropy Genetic Analysis Instrumentation Center for providing the facilities for transcriptional analysis. Special thanks to Dr. Joshua Englert, Emily Shalosky, Dr. John Christman, and Dr. Sangwoon Chung of the Ohio State University for their technical expertise. Additionally, we thank Dr. Fatimah K. Khalaf and BioRender.com for Figure 6A.

